# An Optimized Pipeline for Detection of Salmonella Sequences in Shotgun Metagenomics Datasets

**DOI:** 10.1101/2023.07.27.550528

**Authors:** Lauren M. Bradford, Catherine Carrillo, Alex Wong

## Abstract

**Background:** Culture-independent diagnostic tests (CIDTs) are gaining popularity as tools for detecting pathogens in food. Shotgun sequencing holds substantial promise for food testing as it provides abundant information on microbial communities, but the challenge is in analyzing large and complex sequencing datasets with a high degree of both sensitivity and specificity. Falsely classifying sequencing reads as originating from pathogens can lead to unnecessary food recalls or production shutdowns, while low sensitivity resulting in false negatives could lead to preventable illness.

**Results:** We have developed a bioinformatic pipeline for identifying *Salmonella* as a model pathogen in metagenomic datasets with very high sensitivity and specificity. We tested this pipeline on mock communities of closely related bacteria and with simulated *Salmonella* reads added to published metagenomic datasets. *Salmonella*-derived reads could be found at very low abundances (high sensitivity) without false positives (high specificity). Carefully considering software parameters and database choices is essential to avoiding false positive sample calls. With well-chosen parameters plus additional steps to confirm the taxonomic origin of reads, it is possible to detect pathogens with very high specificity and sensitivity.

## Background

Foodborne illnesses are a global public health issue, with an estimated 600 million incidents of illness and 420 thousand deaths occurring worldwide as of 2010 [1, 2]. In order to prevent consumers from becoming ill, it is essential to detect foodborne pathogens in the production chain.

Culture-based microbiological methods for pathogen detection, which rely on selective enrichment and isolation on agar plates, have been in use for more than a century [3]. Although these methods are sensitive, they are time- and labour-intensive and require labs staffed by expert personnel. In recent years, there has been increasing interest in using culture-independent diagnostic tests (CIDTs) for diagnosis and surveillance of pathogenic organisms of concern. CIDTs include PCR-based methods as well as high-throughput sequencing of either marker genes (e.g 16S rRNA or virulence-related genes) or metagenomes via shotgun sequencing [4, 5, 6, 7, 8, 3, 9].

Shotgun sequencing, wherein all DNA in a sample is sequenced, provides metagenomic data that can be used to detect the presence of pathogens. This type of sequencing avoids the amplification biases that plague phylogenetic metabarcoding [10] and produces datasets containing the full breadth of genetic material [11]. Accordingly, these datasets can also provide information on genes conferring virulence [12] and antimicrobial resistance [3, 13]. Metagenomic data can be used for serotyping of pathogens [14]. It may even be possible to produce metagenome-assembled genomes (MAGs) of pathogens for use in multi-locus sequence typing (MLST) and other analyses [3]. Furthermore, metagenomic datasets can be searched for multiple pathogens during diagnostics or for routine monitoring during food production, although culture enrichment, which requires prior knowledge of possible pathogens-of-interest, is usually still essential to detect organisms at low abundance [3].

While these factors make metagenomics via shotgun sequencing an enticing option for pathogen detection, there are downsides. The pure culture isolates produced by microbiological methods can be used for downstream analyses including drugresistance phenotyping and whole-genome sequencing (WGS) for source attribution [3]; by definition, CIDTs bypass this step [15]. Furthermore, the depth of sequencing required and associated cost must be considered. Detecting low-abundance organisms in samples with overwhelming numbers of reads from the host, food matrix, and/or other microbes is a major barrier [16].

Trustworthy taxonomic classification of each sequencing read is an ongoing challenge, and many bioinformatic tools have been and continue to be developed to address this issue [17, 18, 19, 20, 21, 22, 23]. Metagenomic read classification algorithms primarily rely on identifying species by comparing them to the closest matches in existing databases. However, this approach poses challenges when dealing with species that have limited representation in public repositories, especially when compared to pathogenic species. Additionally, certain sequences exhibit high conservation between species, creating a risk of misclassifying non-pathogens as related pathogens.

Falsely identified reads (that is, sequencing reads erroneously classified as coming from the pathogen of interest) can lead to false positive calls of samples, which presents a particular problem in the field of pathogen detection. In the context of food production, these could cause economic loss from unnecessary recalls or factory shutdowns. Various strategies have been proposed to eliminate false positives, such as setting a high threshold for the number of pathogen-derived reads required for a sample to be considered “positive” [24]; manually curating reference databases and using stringent software settings [25]; or confirming reads putatively classified as the pathogen-of-interest by comparison to species-specific regions (SSRs) [26].

In this study, we aimed to develop and test a pipeline that confidently identifies pathogens from communities of closely related organisms while optimizing lower limits of detection. To do so, we built upon Huang et al.’s [26] use of SSRs to eliminate false positive classification of shotgun sequencing reads. Our interest was particularly in the lower limits of detection, and in confidently identifying pathogens from communities of closely related organisms. We also consider the importance of carefully selecting software parameters (rather than simply using defaults) and reference databases. Kraken [27] and its updated version, Kraken 2 [19] use k-mer based alignment and are among the most highly cited metagenomic classifiers. There are range of pre-made reference databases available for Kraken 2, and it also allows the production of custom databases. With well-chosen databases and parameters, Kraken 2 achieves high precision and recall [28]. For these reasons, Kraken 2 was used as the classifer in this experiment. We use *Salmonella* as a model for the broader problem of pathogen detection in metagenomic datasets. Non-typhoidal serovars of *Salmonella*, which cause potentially life-threatening gastrointestinal illess, are one of the most common contributors to foodborne illness in Canada [29] and are one of the top 4 causes of diarrhoeal diseases globally [30]. However, the findings of this study could be adapted and extended to other pathogens.

## Results

Briefly, our workflow tested reliability and lower limits of detection of simulated *Salmonella* reads in shotgun sequencing datasets. We started with simulated background communities of closely-related bacteria (i.e., members of the Enterobacteriaceae family), since the chance of false identification should be higher with more closely related organisms. We tested classification by Kraken 2 using various reference databases and confidence levels, as well as an additional confirmation step in which putative *Salmonella* reads were compared against “species”-specific reads from the *Salmonella* pan-genome [31]. To compare the detection sensitivity against other reference-based classification software, we also investigated these simulated libraries using the recently-released Metaphlan4 [21].

### Choice of confidence level and database affects number of false positives

We first examined the impact of confidence level. Confidence scoring in Kraken 2 is a simple scheme in which the user defines a score threshold between 0 and 1 (default: 0). Each sequence is scored based on *kmer* mapping, and the label for that sequence is adjusted until the score meets or exceeds the confidence threshold. A more detailed explanation can be found in the software manual^[1]^. At confidence 0, the default setting, the majority of *Salmonella*-derived reads are correctly assigned, but there are many false positives (Fig. 1).

**Figure 1.**
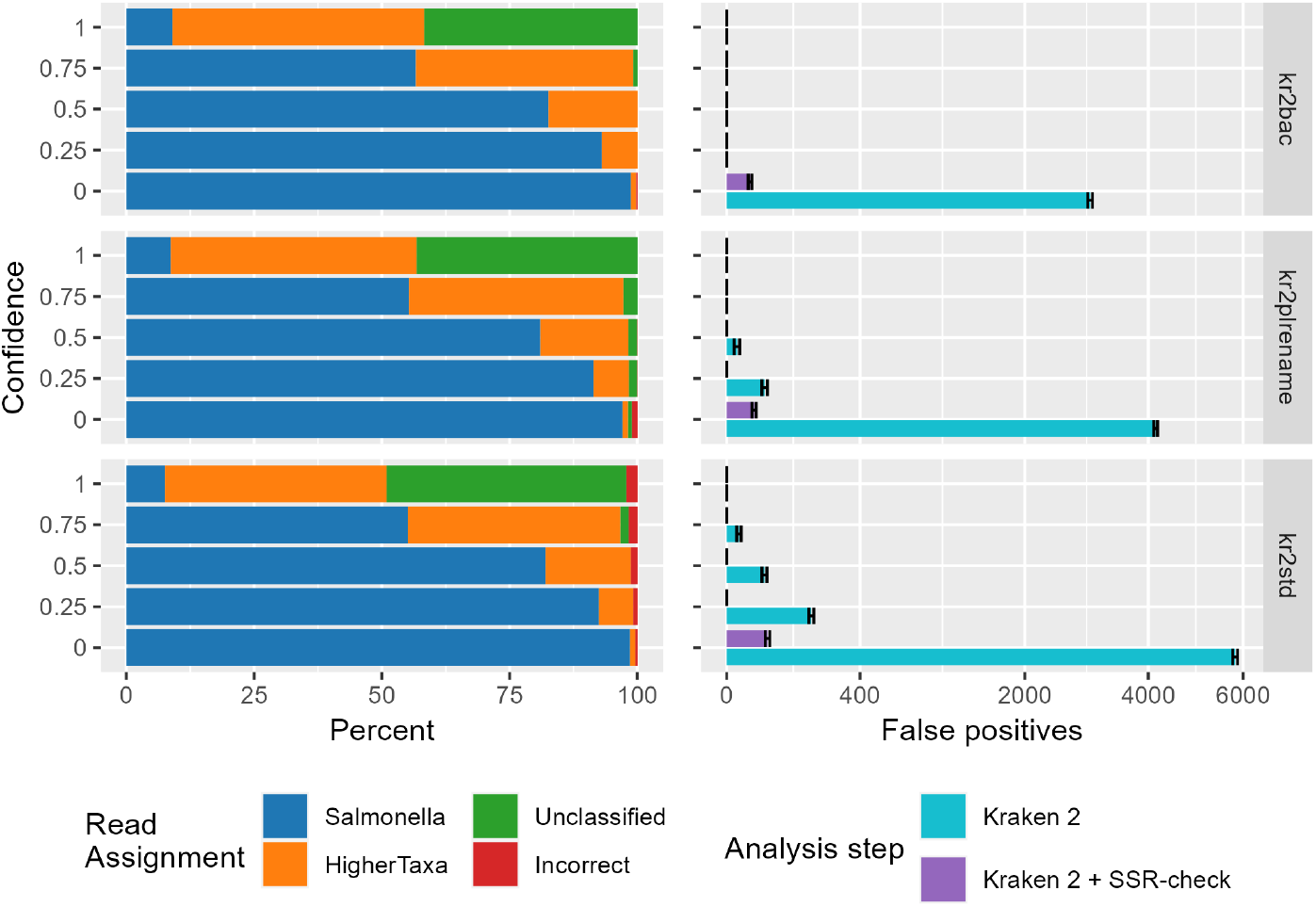
Left panel: Stacked bars showing Kraken 2’s classification of *Salmonella*-derived reads in the library with 0.001% *Salmonella*. In blue, *Salmonella*-derived reads identified explicitly as from the *Salmonella* genus; orange, those identified at a less specific taxonomic level; green, unclassified; and red misidentified as neither *Salmonella* nor an appropriate higher taxonomic group. Right panel: Number of non-*Salmonella*-derived reads classified as *Salmonella* (i.e. false positives) by Kraken 2, and remaining after checking Kraken 2 results against SSRs. Libraries contained 10 million reads each. Error bars are 1 std deviation. X-axis is square-root transformed to better display low values.

As confidence increases, the number of *Salmonella*-derived reads identified higher on the taxonomic tree increases; while these identities are not *incorrect*, they would not lead their libraries to be considered “positive for *Salmonella*”. Strangely, the number of *Salmonella*-derived reads falsely identified (that is, identified as a different genus) increases with increasing confidence when using the standard (“kr2std”) database. Conversely, increasing confidence reduces misidentification when using the bacteria or plasmid-edited bacteria (“kr2plrename”) databases (Fig. 1).

The prevalence of false positives at differing confidence levels can also be seen by their effect on precision in precision-recall curves (Fig. 2). This is most readily apparent in libaries with low counts of *Salmonella*-derived reads (bottom panel). Precision is very low at confidence 0 and near perfect at confidence 1, regardless of database used. However, database choice impacts precision and recall at intermediate confidence levels, with the kr2bac database showing near-perfect precision and high recall already at confidence 0.25 (Fig. 2, bottom panel).

**Figure 2.**
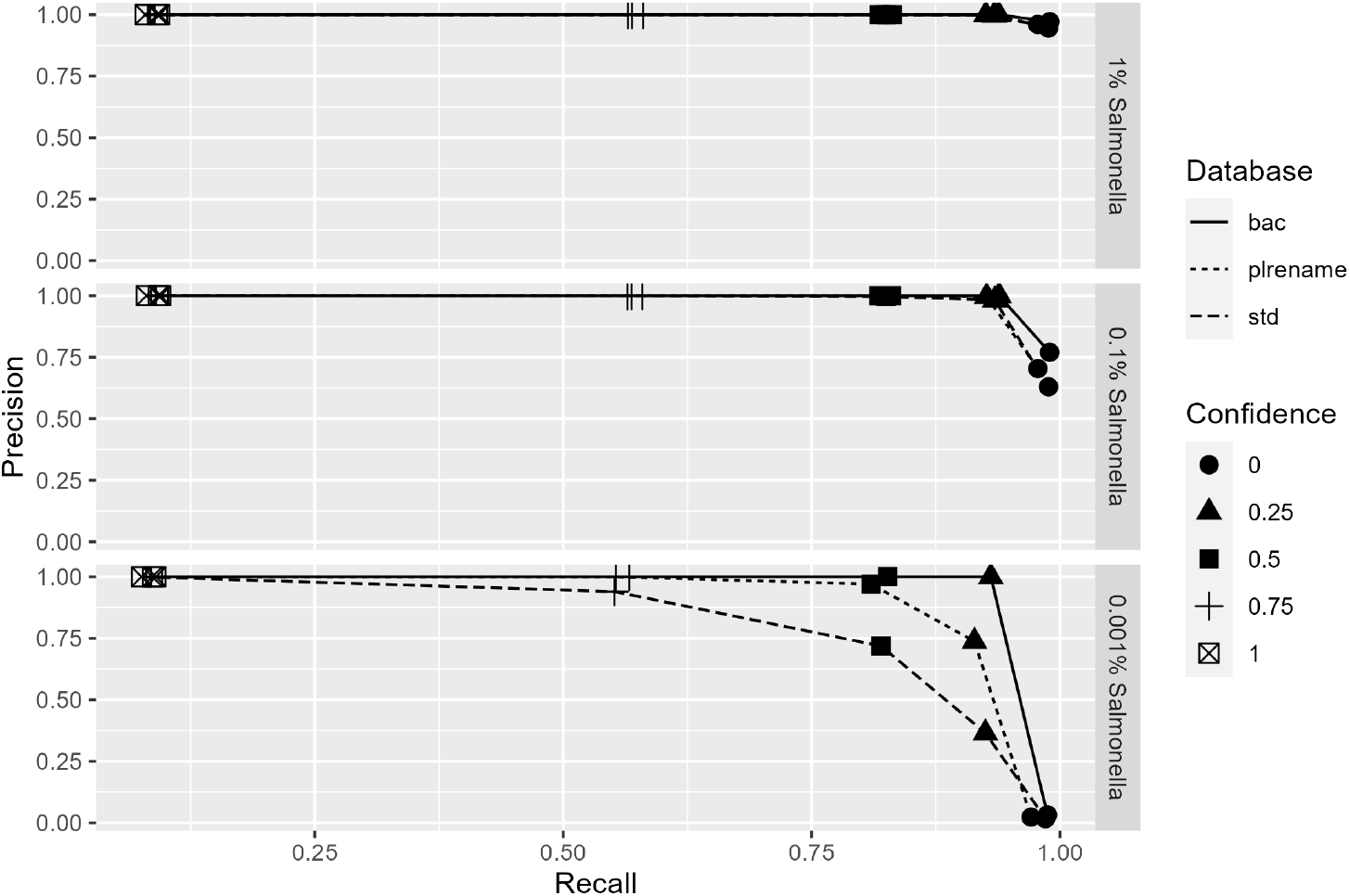
Precision-recall plots for *Salmonella* detection via Kraken 2 classification in 10 million read libraries containing 100k (top panel), 10k (middle panel), and 100 (bottom panel) *Salmonella*-derived reads. Precision is a measure of specificity, with high precision indicating a low rate of false positives; recall is a measure of sensitivity, with high recall indicating a low rate of false negatives.

### Comparison to SSRs is quite effective at removing false positives

To remove false positives while retaining the best chance of detecting true positives, we added a comparison step analogous to that used in the SNIPE pipeline [26]. All reads identified by Kraken 2 as belonging to the *Salmonella* genus were then compared to 403 “species”-specific regions (SSR; though in this case they are genus-rather than species-specific) of 1000 bp length each from the *Salmonella* pan-genome. These SSRs were previously found by Laing et al. [31] by extracting 1000 bp-long regions shared by 211 closed *S. enterica* genomes, iteratively screening these regions against the GenBank nr database, and discarding any region present in any genomic sequence except that of *S. enterica*.

This comparison substantially reduced the number of false positives remaining at the end of the analysis pipeline. For all three databases, however, false positives remained at confidence 0 (the Kraken 2 default) and were only completely absent at confidence *≥* 0.25 (Fig. 1, right panel).

### Reads from novel organisms that are related to Salmonella are also filtered

We have collected genome sequence data for unusual isolates recovered from food and environmental sources, ten of which were mis-identified as *Salmonella* based on closest matches to published genomes by either MASH (1 *Citrobacter* spp.), 16S sequence analysis (6 *Enterobacter/Klebsiella* spp.) or detection of species-specific genes (3 *Citrobacter* spp.). These genomes have not been published and are therefore not incorporated into public databases. To test whether sequencing reads from these organisms pose a problem for the present workflow, a metagenome was created by simulating reads from 31 unpublished genomes to coverage levels of 35 to 82x (Table S2, 10.5281/zenodo.8056523). While Kraken 2 analysis on its own classified 16,904 of the 40 million total reads as coming from *Salmonella*, none of these reads passed through the SSR-check step. Sequencing files are available at 10.5281/zenodo.8056523.

### Workflow limits of detection in a background of related species

A subsequent round of analysis investigated limits of detection using libraries with lower *Salmonella* content and the analysis parameters with the best precision and recall characteristics, i.e., the Kraken 2 bacteria database and confidence 0.25, with subsequent confirmation of putative *Salmonella* reads by comparison to SSRs. All libraries contained 10 million total reads, and included 5, 10, or 50 *Salmonella*-derived reads. At least one read was positively identified as *Salmonella* in 16/20 replicates of 50 *Salmonella* read libraries, 14/20 replicates of 10 *Salmonella* read libraries, and 12/20 replicates of 5 *Salmonella* read libraries (Table 1), giving a calculated LOD_50_ of 10.2 reads in a 10 million read library [32] (CI: 6.8-15.3). In comparison, Metaphlan4 was much less sensitive, requiring 1 *×* 10^4^ *Salmonella*-derived reads in a 10 million read library (0.1 %) for reliable detection (Table 1), with a calculated LOD_50_ of 2106 reads in a 10 million read library (CI: 1247-3557).

**Table 1.** Number of replicate libraries at each spike-in level testing positive for *Salmonella* according to the current pipeline (“Kraken2 + SSR”) or Metaphlan4. Libraries contained 10 million reads.

| Salmonella reads<br>in library | Replicates | Positive libraries:<br>Kraken2+SSRs | Positive libraries:<br>Metaphlan4 |
| --- | --- | --- | --- |
| $1 \times 10^5$ | 20 | 20 | 20 |
| $1 \times 10^4$ | 20 | 20 | 20 |
| $1 \times 10^3$ | 20 | 20 | 4 |
| $1 \times 10^2$ | 20 | 20 | 0 |
| 50 | 20 | 16 | 0 |
| 10 | 20 | 13 | 0 |
| 5 | 20 | 12 | 0 |
| 0 | 20 | 0 | 0 |

### Limits of detection in a real microbiome background

Previous rounds of analysis made use of fully simulated shotgun sequencing libraries containing reads from members of the Enterobacteriaceae family. To explore use of the analysis pipeline for detecting *Salmonella* in more realistic set of sequences, libraries were created using published shotgun sequencing datasets from chicken gut microbiomes. *Salmonella* detection was attempted by (a) searching for two *Salmonella* marker genes and (b) using the pipeline established above (using the Kraken 2 bacteria database and confidence 0.25, plus comparison to *Salmonella* SSRs). Marker genes *invA* and *stn* are commonly used for *Salmonella* detection in rapid tests such as quantitative PCR and loop-mediated isothermal amplification [33, 34, 35]. Read fragments of these genes could be reliably detected (100 % of replicates) in libraries with approx. 4*×*10^4^ *Salmonella*-derived reads, with an LOD_50_ for one or more of the markers of 1754 (CI: 1067-2884) *Salmonella*-derived reads in a 40 million read library (Fig. 3, top panel). Using the established detection pipeline, *Salmonella* reads could be detected in all libraries with 40 *Salmonella*-derived reads, with an LOD_50_ of 5.5 (CI: 3.1-9.8).

**Figure 3.**
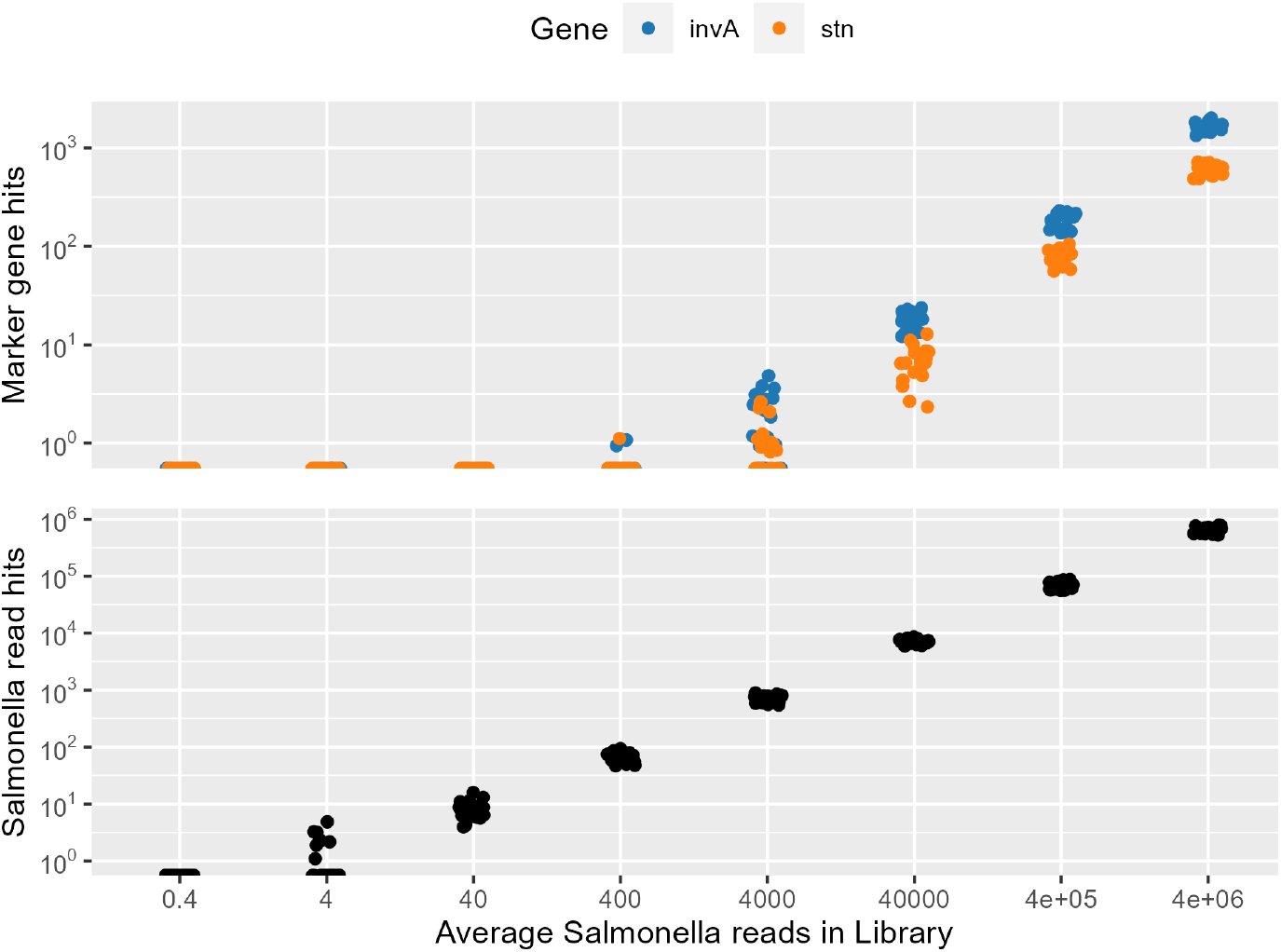
Detection of *Salmonella* marker genes (top) or *Salmonella*-derived reads (bottom) using the established workflow in a chicken caecal microbiome background. Libraries contained 40 million total reads. Datapoints from individual replicates are shown. Y-axis is in log10 scale.

## Discussion

Here we present a metagenomics analysis pipeline for the confident detection of *Salmonella* in a background matrix containing closely related species, which could pose problems in the form of false positives. We show the importance of appropriate database and software parameter choices, and extend the SNIPE pipeline’s [26] use of species-specific regions (SSRs) for identifying and filtering false positives given by popular classification software Kraken 2 [19]. In establishing their pipeline, Huang et al. used default parameters in their chosen classification software, which this current study has found to be inadequate. At confidence 0 (the Kraken 2 default), false positives persist even after the SSR-comparison filtering step. Furthermore, their testing sets included a very limited number of closely related genomes as a confounding factor, whereas multiple members of the Enterobacteriaceae family could be expected to be present in sample types that are frequent targets for pathogen detection, including human clinical samples [36], food-animal microbiomes [37], or food products [38]. Thus, testing extensively in a dataset containing a large number and variety of related organisms was informative.

We were particularly interested in the sensitivity of pathogen detection. Low limits of detection make it possible to detect *Salmonella* even when it is a very small component of the sample community. Additionally, extremely sensitive bioinformatic methods allow detection from shallower sequencing datasets, which would reduce costs. Our pipeline was able to correctly identify 100 % of *Salmonella*-positive sequencing libraries containing just 100 Salmonella-derived reads. Even with just five Salmonella-derived reads, more than half of library replicates were correctly identified as positive. By comparison, the recently-released Metaphlan4 software was very specific, but far less sensitive. However, one consideration in using such a sensitive detection pipeline is the risk contamination via carry-over between sequencing runs, a known issue with Illumina sequencers [39]. Samples contaminated in this way would legitimately contain reads identified as belonging to the pathogen of interest by this pipeline, and thus be considered positive [24, 40]. There is presently no way to overcome this issue in data analysis once sequencing has been performed; it can only be minimized during wet-lab procedures.

There are additional limitations to this analysis. Almost all members of the *Salmonella* genus are considered pathogenic [41], so identification at the genus level is sufficient for these organisms. Other genera contain both benign and pathogenic members, making species-level identification necessary. Still other species or sub-species are benign unless they carry certain virulence factors (for example, the majority of *E. coli* are harmless, but Shiga-toxin producing *E. coli* (STEC) cause gastrointestinal illness and even death [42]). In such cases, virulence genes or genetically linked markers must be detected for positive identification [15, 43]. We show that far higher pathogen numbers in a population are required for detection of marker genes (in this study, *invA* and *stn*) compared to general genomic reads.

We found that best results came from using the Kraken 2 bacteria database; however, we had prior knowledge that the pathogen-of-interest is bacterial. Diagnostic analyses where the cause of disease is unknown would require use of additional databases (ex. a virus database), and many additional SSRs for various species. The kraken 2-build function allows the production of custom databases, so it would be possible to create a combined bacteria-virus database, and to add in organisms of interest that are not yet included.

## Conclusions

Shotgun sequencing is gaining popularity in many biological fields, including food safety. However, it is challenging to analyze the resulting datasets for the presence of pathogens with a high degree of both sensitivity and specificity. Believable results as far as positive and negative calls for samples are essential when false positives could lead to food recalls or production shut-downs and false negatives could lead to preventable illnesses. Many pipelines exist for metagenomics-based detection of foodborne pathogens [24], but these pipelines are often not tested on mock communities where the provenance of each read is known and false classification can be assessed. Here, we have built upon previous suggestions to develop and systematically test a pipeline for detection of *Salmonella* as a model pathogen in metagenomic datasets. We emphasize that careful consideration of software parameter and database choices is essential. With well-chosen parameters plus additional steps to confirm the taxonomic origin of reads, it is possible to detect pathogens with very high specificity and sensitivity.

## Methods

### Mock Community

The mock community “enterobac” is composed of members of the Enterobaceriaceae family, to which the genus *Salmonella* belongs. Complete reference genomes for 62 species in the Enterobacteraceae family were selected using the NCBI genome browser^[2]^ and downloaded from the NCBI RefSeq database (Table S1).

The art illumina function of ART [44] was used to generate simulated shotgun sequencing reads for each genome with the following parameters: 25-fold coverage, paired reads of length 150 bp with insert size 300 bp, read length standard deviation of 10 bp, and an error profile from the Illumina HiSeq 2500. Reads from all genomes except *Salmonella enterica* subsp. enterica serovar Typhimurium str. LT2 were concatenated into “master” mock community files with a total of 26,976,269 paired-end reads.

### Mock Libraries

Libraries of 10 million paired reads were produced by randomly subsetting reads from the master file using BBMAPs’ reformat function [45]. Read counts were chosen based on the desired number of *Salmonella* reads per library; for example, for the 10 % *Salmonella* library, 1 *×* 10^5^ reads were selected from those produced from the *Salmonella* Typhimurium genome, and 9 *×* 10^5^ reads were selected from the master mock community file. Twenty replicates were produced at each target level.

An additional mock community was also generated, comprised of unpublished genomic data from 31 strains erroneously identified as *Salmonella* by either MASH (1 *Citrobacter* spp.), 16S sequence analysis (6 *Enterobacter/Klebsiella* spp.) or detection of species-specific genes (3 *Citrobacter* spp.) (Table S2). Genomes had 35 to 82 fold coverage, with a total of 40 Million paired end reads. Libraries were produced similar to above. Illumina HiSeq short reads were synthesized from the draft genome assemblies and raw reads of the bacterial genomes using the FetaGenome2 (fabricate metagenome) tool developed in house^[3]^. Briefly, Art version 2.5.8 was used to simulate paired-end HiSeq reads of 150 bp in length with a 300 bp insert size. To simulate variability in coverage levels (e.g. higher coverage in plasmids vs chromosomal sequences), the FetaGenomePlasmidAware edition uses BWA to map reads to the original assembly to determine coverage depth of each contig in the given assembly, then uses the coverage report output to create more reads for higher-depth locations and fewer reads for low-depth locations of the genome. The simulated library was tested with the current workflow, with Kraken 2 confidence of 0.25 and the kr2bac database, followed by confirmation by checking against Salmonella SSRs.

### Kraken 2 reference databases

A pre-indexed version of the Kraken 2 [19] standard database (“kr2std”), which contains archaea, bacteria, viral, plasmid, human, and UniVec Core sequences^[4]^ was downloaded on 01 Oct 2021 (database last updated 17 May 2021).

The Kraken 2 bacteria library and taxonomy were downloaded on 28 Oct 2021 according to the software manual instructions (see supplementary material). The unaltered Kraken 2 bacteria databases (“kr2bac”) was built using these files. Database “kr plrenamed db” was built after altering the bacteria library file according to instructions from Doster et al. (2019) [25] (see supplementary material). Plasmids in the bacteria library fasta file were renamed using sed, and the database was then built as above.

### *Salmonella* Species Specific Regions (SSRs)

Laing et al. (2017) [31] investigated the *Salmonella* pan-genome and found 403 regions of 1000 bp each that were specific to the *Salmonella* genus. These regions were used to confirm the identity of reads classified as *Salmonella*-derived by Kraken 2. The position of on these regions on the *Salmonella* reference genome (*Salmonella enterica* subsp. enterica serovar Typhimurium str. LT2) was taken from the supplementary files [31] and the faidx function of samtools [46] was used to extract the sequences in fasta format. A blast-formatted database was then created using the sequences and BLAST CLI’s makeblastdb command [47, 48].

### Workflows

Custom Snakemake [49] workflows were written to carry out library setup and analyses. Each mock library was first subject to trimming with Trimmomatic [50] with parameters SLIDINGWINDOW:4:20 MINLEN:36. Singleton files passing quality check from the Trimmomatic output were concatenated with paired files to ensure minimal loss of sequences. Reads were then classified with Kraken 2 [19]. The first round of analysis was used to establish the best database and confidence level. For this, 10 million-read libraries with 1 %, 0.1 %, and 0.01 % *Salmonella* content were classified with each of the three Kraken 2 databases described above (kr2std, kr2bac, and kr2plrename) at five confidence levels: 0 (default), 0.25, 0.5, 0.75, and 1.

Output from this analysis was also used to establish the utility of comparison to SSRs [31] for removing false positives. Information about reads classified as members of the *Salmonella* genus (“putative *Salmonella* hits”) was extracted from the Kraken 2 output and the origin of the read recorded using a custom Python script to determine the number of false positives (that is, reads originating from a non-*Salmonella* genome that were classified as a member of the *Salmonella* genus). Sequences from all Kraken 2 *Salmonella* hits were compared to the SSR database using the BLAST command line application [47]. The origin of these SSR *Salmonella* hits was again checked to determine remaining false positives.

The second round of analysis explored lower limits of detection in mock communities based on best practices from the above analyses. Libraries with 0.005 %, 0.001 %, and 0.0005 % *Salmonella* were classified with Kraken 2 against the bacteria database (“kr2bac”) at 0.25 confidence. Kraken 2 *Salmonella* hits were extracted and compared to SSRs, and false positives were recorded, as above.

Mock libraries were also analyzed with Metaphlan4 [21] using the vJan21 database. All reads that passed the Trimmomatic step were combined into one file per library and analyzed with default parameters, using the output parameters “unclassified estimation” and “-t rel ab w read stats”. Individual library profiles were combined with the merge metaphlan tables.py script, and libraries with at least one read in the *Salmonella* genus were considered positive for *Salmonella*.

### Limits of detection in real metagenomic background

Limits of detection for *Salmonella*-derived reads were further explored using published chicken caecal shotgun libraries as the background microbiome. Sequencing files from Salaheen et al. (2017) [51] were retrieved from the European Nucleotide Archive (accession codes SRR5280289, SRR5280393, and SRR5280514). Briefly, the Salaheen et al. (2017) study investigated the impact of antibiotic growth promotors on cecal microbiomes of Cobb-500 broiler chicks. Retrieved sequences were from control chickens which did not receive growth promotors. These reads were paired-end (2×151 bp) from an Illumina NextSeq 500. Sequencing files were concatenated to create master microbiome files containing 119,068,070 paired reads. The master files were classified with Kraken 2 [19] using the kr2bac library and confidence 0.2, and all reads matching to *Salmonella* were removed. This resulted in master files of 119,068,030 reads. Reads were also checked against a custom database derived from the chicken (Gallus gallus) reference genome to ensure that the number of host reads in the shotgun dataset was neglible.

The genome of *Salmonella* Enteritidis strain CFIAFB20140150 (accession code SRR10859048) [52] was used for generation of simulated shotgun sequencing reads using the art illumina function of ART [44], as above. This strain was chosen based on its concurrent use in a laboratory spike-in study. Replicate libraries were produced by appending the appropriate number of *Salmonella* Enteritidis-derived reads to the master microbiome files, then subsetting libraries of 40 million reads using BBMAP’s reformat function [45]. Libraries were produced at eight target levels, from 10 % (approx. 4 million S. Enteritidis-derived reads) to 0.000001 % (0.4 reads), and 20 replicates were produced per target level.

Libraries were analyzed using the above workflow, with the Kraken 2 bacteria database, confidence 0.25, and SSR checks. Additionally, DIAMOND-formatted databases of the *invA* and *stn* marker genes were created using animo acid sequences retrieved from NCBI (WP 000927219.1 and AAA21354.1, respectively) with the DI-AMOND makedb function [53]. The presence of these genes in libraries was tested using DIAMOND’s blastx function with a percent ID cutoff of 96.

### Statistics

Plotting and statistical analyses were carried out in R v4.2.2 [54]. The full list of packages used is available in the supplementary material. LOD_50_ was calcluated via the log-log model by Wilrich and Wilrich [32] using a tool they provide online^[5]^. Although this model was developed for calculating LODs in terms of bacterial CFU per gram of food matrix during spike-in experiments, we adapted the calculation for counts of pathogen-derived reads in sequencing libraries.

## Acknowledgements

We are grateful for funding from the Ontario Ministry of Agriculture, Food, and Rural Affairs (OMAFRA). This research was enabled in part by support provided by SHARCNET (sharcnet.ca) and the Digital Research Alliance of Canada (alliancecan.ca). Thanks to Ashley Cooper at the CFIA for producing the mock library from unpublished genomes, and to Ryan Taylor at Carleton University for sharing his technical expertise.

## Availability of data and materials

Genome sequences of unusual isolates are available at 10.5281/zenodo.8056523. Analysis code can be viewed at https://github.com/LMBradford/salmdetectpipeline.git.

## Authors’ contributions

LMB conceived and carried out the study. LMB also drafted the manuscript. AW and CC contributed resources and revised the manuscript.

## Footnotes

[1] https://github.com/DerrickWood/kraken2/wiki/Manual

[2] https://www.ncbi.nlm.nih.gov/datasets/genomes/?taxon=543

[3] https://github.com/OLC-Bioinformatics/FetaGenome2

[4] https://benlangmead.github.io/aws-indexes/k2

[5] https://www.wiwiss.fu-berlin.de/fachbereich/vwl/iso/ehemalige/wilrich/index.html

## References

1. World Health Organization: WHO Estimates of the Global Burden of Foodborne Diseases: Foodborne Disease Burden Epidemiology Reference Group 2007-2015. World Health Organization, Geneva (2015)

2. Lee, H., Yoon, Y.: Etiological agents implicated in foodborne illness world wide. Food science of animal resources 41(1), 1 (2021)

3. Banerjee, G., Agarwal, S., Marshall, A., Jones, D.H., Sulaiman, I.M., Sur, S., Banerjee, P.: Application of advanced genomic tools in food safety rapid diagnostics: challenges and opportunities. Current Opinion in Food Science, 100886 (2022)

4. Koutsoumanis, K., Allende, A., Alvarez-Ordóñez, A., Bolton, D., Bover-Cid, S., Chemaly, M., Davies, R., De Cesare, A., Hilbert, F., et al.: Whole genome sequencing and metagenomics for outbreak investigation, source attribution and risk assessment of food-borne microorganisms. EFSA Journal 17(12), 05898 (2019)

5. Bell, R.L., Jarvis, K.G., Ottesen, A.R., McFarland, M.A., Brown, E.W.: Recent and emerging innovations in salmonella detection: a food and environmental perspective. Microbial biotechnology 9(3), 279–292 (2016)

6. Muhamad Rizal, N.S., Neoh, H.-m., Ramli, R., A/LK Periyasamy, P.R., Hanafiah, A., Abdul Samat, M.N., Tan, T.L., Wong, K.K., Nathan, S., Chieng, S., et al.: Advantages and limitations of 16s rrna next-generation sequencing for pathogen identification in the diagnostic microbiology laboratory: perspectives from a middle-income country. Diagnostics 10(10), 816 (2020)

7. Yap, M., Ercolini, D., Álvarez-Ordóñez, A., O’Toole, P.W., O’Sullivan, O., Cotter, P.D.: Next-generation food research: use of meta-omic approaches for characterizing microbial communities along the food chain. Annual Review of Food Science and Technology 13, 361–384 (2022)

8. Jagadeesan, B., Gerner-Smidt, P., Allard, M.W., Leuillet, S., Winkler, A., Xiao, Y., Chaffron, S., Van Der Vossen, J., Tang, S., Katase, M., et al.: The use of next generation sequencing for improving food safety: translation into practice. Food microbiology 79, 96–115 (2019)

9. Forbes, J.D., Knox, N.C., Ronholm, J., Pagotto, F., Reimer, A.: Metagenomics: the next culture-independent game changer. Frontiers in microbiology 8, 1069 (2017)

10. Shah, N., Tang, H., Doak, T.G., Ye, Y.: Comparing bacterial communities inferred from 16s rrna gene sequencing and shotgun metagenomics. In: Biocomputing 2011, pp. 165–176. World Scientific, Singapore (2011)

11. Ranjan, R., Rani, A., Metwally, A., McGee, H.S., Perkins, D.L.: Analysis of the microbiome: Advantages of whole genome shotgun versus 16s amplicon sequencing. Biochemical and biophysical research communications 469(4), 967–977 (2016)

12. Yang, X., Noyes, N.R., Doster, E., Martin, J.N., Linke, L.M., Magnuson, R.J., Yang, H., Geornaras, I., Woerner, D.R., Jones, K.L., et al.: Use of metagenomic shotgun sequencing technology to detect foodborne pathogens within the microbiome of the beef production chain. Applied and environmental microbiology 82(8), 2433–2443 (2016)

13. Duarte, A.S.R., Röder, T., Van Gompel, L., Petersen, T.N., Hansen, R.B., Hansen, I.M., Bossers, A., Aarestrup, F.M., Wagenaar, J.A., Hald, T.: Metagenomics-based approach to source-attribution of antimicrobial resistance determinants–identification of reservoir resistome signatures. Frontiers in microbiology 11, 601407 (2021)

14. Zhang, S., Yin, Y., Jones, M.B., Zhang, Z., Deatherage Kaiser, B.L., Dinsmore, B.A., Fitzgerald, C., Fields, P.I., Deng, X.: Salmonella serotype determination utilizing high-throughput genome sequencing data. Journal of clinical microbiology 53(5), 1685–1692 (2015)

15. Carleton, H.A., Besser, J., Williams-Newkirk, A.J., Huang, A., Trees, E., Gerner-Smidt, P.: Metagenomic approaches for public health surveillance of foodborne infections: opportunities and challenges. Foodborne Pathogens and Disease 16(7), 474–479 (2019)

16. Cocolin, L., Mataragas, M., Bourdichon, F., Doulgeraki, A., Pilet, M.-F., Jagadeesan, B., Rantsiou, K., Phister, T.: Next generation microbiological risk assessment meta-omics: the next need for integration. International Journal of Food Microbiology 287, 10–17 (2018)

17. Sczyrba, A., Hofmann, P., Belmann, P., Koslicki, D., Janssen, S., Dröge, J., Gregor, I., Majda, S., Fiedler, J., Dahms, E., et al.: Critical assessment of metagenome interpretation—a benchmark of metagenomics software. Nature methods 14(11), 1063–1071 (2017)

18. Meyer, F., Fritz, A., Deng, Z.-L., Koslicki, D., Lesker, T.R., Gurevich, A., Robertson, G., Alser, M., Antipov, D., Beghini, F., et al.: Critical assessment of metagenome interpretation: the second round of challenges. Nature methods 19(4), 429–440 (2022)

19. Wood, D.E., Lu, J., Langmead, B.: Improved metagenomic analysis with kraken 2. Genome biology 20(1), 1–13 (2019)

20. Huson, D.H., Beier, S., Flade, I., Górska, A., El-Hadidi, M., Mitra, S., Ruscheweyh, H.-J., Tappu, R.: Megan community edition-interactive exploration and analysis of large-scale microbiome sequencing data. PLoS computational biology 12(6), 1004957 (2016)

21. Blanco-Míguez, A., Beghini, F., Cumbo, F., McIver, L.J., Thompson, K.N., Zolfo, M., Manghi, P., Dubois, L., Huang, K.D., Thomas, A.M., et al.: Extending and improving metagenomic taxonomic profiling with uncharacterized species using metaphlan 4. Nature Biotechnology, 1–12 (2023)

22. Menzel, P., Ng, K.L., Krogh, A.: Fast and sensitive taxonomic classification for metagenomics with kaiju. Nature communications 7(1), 1–9 (2016)

23. Ounit, R., Wanamaker, S., Close, T.J., Lonardi, S.: Clark: fast and accurate classification of metagenomic and genomic sequences using discriminative k-mers. BMC genomics 16(1), 1–13 (2015)

24. Höper, D., Grützke, J., Brinkmann, A., Mossong, J., Matamoros, S., Ellis, R.J., Deneke, C., Tausch, S.H., Cuesta, I., Monzón, S., et al.: Proficiency testing of metagenomics-based detection of food-borne pathogens using a complex artificial sequencing dataset. Frontiers in microbiology 11, 575377 (2020)

25. Doster, E., Rovira, P., Noyes, N.R., Burgess, B.A., Yang, X., Weinroth, M.D., Linke, L., Magnuson, R., Boucher, C., Belk, K.E., et al.: A cautionary report for pathogen identification using shotgun metagenomics; a comparison to aerobic culture and polymerase chain reaction for salmonella enterica identification. Frontiers in microbiology 10, 2499 (2019)

26. Huang, L., Hong, B., Yang, W., Wang, L., Yu, R.: Snipe: highly sensitive pathogen detection from metagenomic sequencing data. Briefings in Bioinformatics 22(5), 064 (2021)

27. Wood, D.E., Salzberg, S.L.: Kraken: ultrafast metagenomic sequence classification using exact alignments. Genome biology 15(3), 1–12 (2014)

28. Wright, R.J., Comeau, A.M., Langille, M.G.: From defaults to databases: parameter and database choice dramatically impact the performance of metagenomic taxonomic classification tools. bioRxiv (2022)

29. Thomas, M.K., Murray, R., Flockhart, L., Pintar, K., Pollari, F., Fazil, A., Nesbitt, A., Marshall, B.: Estimates of the burden of foodborne illness in canada for 30 specified pathogens and unspecified agents, circa 2006. Foodborne pathogens and disease 10(7), 639–648 (2013)

30. World Health Organization: Salmonella (non-typhoidal) [Fact Sheet]. https://www.who.int/news-room/fact-sheets/detail/salmonella-(non-typhoidal)

31. Laing, C.R., Whiteside, M.D., Gannon, V.P.: Pan-genome analyses of the species salmonella enterica, and identification of genomic markers predictive for species, subspecies, and serovar. Frontiers in microbiology 8, 1345 (2017)

32. Wilrich, C., Wilrich, P.-T.: Estimation of the pod function and the lod of a qualitative microbiological measurement method. Journal of AOAC International 92(6), 1763–1772 (2009)

33. Rahn, K., De Grandis, S., Clarke, R., McEwen, S., Galan, J., Ginocchio, C., Curtiss Iii, R., Gyles, C.: Amplification of an inva gene sequence of salmonella typhimurium by polymerase chain reaction as a specific method of detection of salmonella. Molecular and cellular probes 6(4), 271–279 (1992)

34. Moore, M., Feist, M.: Real-time pcr method for salmonella spp. targeting the stn gene. Journal of applied microbiology 102(2), 516–530 (2007)

35. Ou, H., Wang, Y., Gao, J., Bai, J., Zhang, Q., Shi, L., Wang, X., Wang, C.: Rapid detection of salmonella based on loop-mediated isothermal amplification. Annals of palliative medicine 10(6), 6850–6858 (2021)

36. King, C.H., Desai, H., Sylvetsky, A.C., LoTempio, J., Ayanyan, S., Carrie, J., Crandall, K.A., Fochtman, B.C., Gasparyan, L., Gulzar, N., et al.: Baseline human gut microbiota profile in healthy people and standard reporting template. PloS one 14(9), 0206484 (2019)

37. Shang, Y., Kumar, S., Oakley, B., Kim, W.K.: Chicken gut microbiota: importance and detection technology. Frontiers in Veterinary Science 5, 254 (2018)

38. Baylis, C., Uyttendaele, M., Joosten, H., Davies, A., Heinz, H.: The enterobacteriaceae and their significance to the food industry, ilsi europe report series. Technical report, Washington, DC: International Life Sciences Institute (2011)

39. Illumina: Reducing run-to-run carryover on the MiSeq using dilute sodium hypochlorite solution. San Diego, CA Illumina (2013)

40. Nelson, M.C., Morrison, H.G., Benjamino, J., Grim, S.L., Graf, J.: Analysis, optimization and verification of illumina-generated 16s rrna gene amplicon surveys. PloS one 9(4), 94249 (2014)

41. Eng, S.-K., Pusparajah, P., Ab Mutalib, N.-S., Ser, H.-L., Chan, K.-G., Lee, L.-H.: Salmonella: a review on pathogenesis, epidemiology and antibiotic resistance. Frontiers in Life Science 8(3), 284–293 (2015)

42. Majowicz, S.E., Scallan, E., Jones-Bitton, A., Sargeant, J.M., Stapleton, J., Angulo, F.J., Yeung, D.H., Kirk, M.D.: Global incidence of human shiga toxin–producing escherichia coli infections and deaths: a systematic review and knowledge synthesis. Foodborne pathogens and disease 11(6), 447–455 (2014)

43. Riley, L.W.: Distinguishing pathovars from nonpathovars: Escherichia coli. Microbiology Spectrum 8(4), 8–4 (2020)

44. Huang, W., Li, L., Myers, J.R., Marth, G.T.: Art: a next-generation sequencing read simulator. Bioinformatics 28(4), 593–594 (2012)

45. Bushnell, B.: Bbmap: a fast, accurate, splice-aware aligner. Technical report, Lawrence Berkeley National Lab.(LBNL), Berkeley, CA (United States) (2014)

46. Li, H., Handsaker, B., Wysoker, A., Fennell, T., Ruan, J., Homer, N., Marth, G., Abecasis, G., Durbin, R.: The sequence alignment/map format and samtools. Bioinformatics 25(16), 2078–2079 (2009)

47. National Center for Biotechnology Information (US): BLAST Command Line Applications User Manual. (2008)

48. Madden, T.: The blast sequence analysis tool. The NCBI handbook (2003)

49. Mölder, F., Jablonski, K.P., Letcher, B., Hall, M.B., Tomkins-Tinch, C.H., Sochat, V., Forster, J., Lee, S., Twardziok, S.O., Kanitz, A., et al.: Sustainable data analysis with snakemake. F1000Research 10 (2021)

50. Bolger, A.M., Lohse, M., Usadel, B.: Trimmomatic: a flexible trimmer for illumina sequence data. Bioinformatics 30(15), 2114–2120 (2014)

51. Salaheen, S., Kim, S.-W., Haley, B.J., Van Kessel, J.A.S., Biswas, D.: Alternative growth promoters modulate broiler gut microbiome and enhance body weight gain. Frontiers in microbiology 8, 2088 (2017)

52. Cooper, A.L., Low, A.J., Koziol, A.G., Thomas, M.C., Leclair, D., Tamber, S., Wong, A., Blais, B.W., Carrillo, C.D.: Systematic evaluation of whole genome sequence-based predictions of salmonella serotype and antimicrobial resistance. Frontiers in microbiology 11, 549 (2020)

53. Buchfink, B., Reuter, K., Drost, H.-G.: Sensitive protein alignments at tree-of-life scale using diamond. Nature methods 18(4), 366–368 (2021)

54. R Core Team: R: A Language and Environment for Statistical Computing. R Foundation for Statistical Computing, Vienna, Austria (2021). R Foundation for Statistical Computing. https://www.R-project.org/

