## Supplementary information for "An Optimized Pipeline for Detection of Salmonella Sequences in Shotgun Metagenomics Datasets"

**Kraken 2 database commands**

### Kraken 2 bacteria library

```
kraken2-build --download-library bacteria --db kr\_bac\_db

kraken2-build --download-taxonomy --db kr\_bac\_db

kraken2-build --build --db kr\_bac\_db --threads 32
```

#Kraken 2 plrenamed library

Reference 1 suggested that in order to increase specificity and account for horizontal gene transfer, all plasmids in a Kraken 2 database should be given the NCBI taxid 45202 ("unidentified plasmid"), rather than being associated with their taxonomy-of-origin. In their study, this action decreased the number of samples considered *Salmonella*-positive by Kraken 2 analysis and increased the concordance between shotgun sequencing and qPCR/aerobic culture results. Renaming was accomplished after library download by concatenating fasta files and editing plasmid names with a custom sed command:

```
cat $DBNAME/library/*/library.fna > $DBNAME/library/concatenated_genomes.fna
sed -i '/plasmid/c\>kraken:taxid|45202|plasmid' concatenated_library_genomes.fna
```

Full instructions can be found on their github site: https://github.com/colostatemeg/meglab-kraken-custom-db .

**Unpublished genomes mock community**

Illumina HiSeq short reads were synthesized from the draft genome assemblies and raw reads of the bacterial genomes using the FetaGenome2 (fabricate metagenome) tool developed in house (available at: https://github.com/OLC-Bioinformatics/FetaGenome2).

```
FetaGenomePlasmidAware -c FetaConfig -f Synthetic_metagenome -n 40000000 -l 150 -i
300 -t 10 -p HS25
```

**Data analysis packages**

Outputs from the snakemake workflow were summarized with custom Python scripts using pandas [2, 3].

Plotting and statistical analyses were carried out in R v4.2.2 [4] using packages egg [5], dplyr [6], ggplot2 [7], reshape2 [8], tidyr [9], and the ggsci colour palette [10].

Table S1 (provided as .xlsx): Genomes used in the production of the *enterobac* mock community

Table S2 (provided as .xlsx): Genomes used in the production of the additional mock community (previously unpublished organisms)

#### SI References


1. Doster E, Rovira P, Noyes NR, Burgess BA, Yang X, Weinroth MD, Linke L, Magnuson R, Boucher C, Belk KE, others: **A cautionary report for pathogen identification using shotgun metagenomics; a comparison to aerobic culture and polymerase chain reaction for Salmonella enterica identification**. *Frontiers in microbiology* 2019, **10**:2499.

2. Pandas development team T: **pandas-dev/pandas: Pandas**. 2020.

3. McKinney W: **Data Structures for Statistical Computing in Python** . In *Proceedings of the 9th Python in Science Conference* . edited by Stéfan van der Walt and Jarrod Millman 2010: 56 – 61 .

4. Team RC: *R: A Language and Environment for Statistical Computing*. Vienna, Austria: 2021.

5. Auguie B: *egg: Extensions for "ggplot2": Custom Geom, Custom Themes, Plot Alignment, Labelled Panels, Symmetric Scales, and Fixed Panel Size*. 2019.

6. Wickham H, François R, Henry L, Müller K: *dplyr: A Grammar of Data Manipulation*. 2022.

7. Wickham H: *ggplot2: Elegant Graphics for Data Analysis*. Springer-Verlag New York; 2016.

8. Wickham H: **Reshaping Data with the reshape Package**. *Journal of Statistical Software* 2007, **21**:1–20.

9. Wickham H, Girlich M: *tidyr: Tidy Messy Data*. 2022.

10. Xiao N: *ggsci: Scientific Journal and Sci-Fi Themed Color Palettes for "ggplot2."*2023.
